# Bacterial defenses and their trade-off with growth are not ubiquitous but depend on ecological contexts

**DOI:** 10.1101/2024.03.24.586452

**Authors:** Zhi-Ling Liu, Jia Liu, Deng-Ke Niu

## Abstract

Bacteriophages, also known as bacterial viruses, significantly influence microbial ecosystems, driving bacteria to evolve diverse antiviral defense mechanisms. This study explores the intricate relationship between bacterial defenses and growth rates across diverse ecological contexts. Our investigation reveals that bacteria lacking defenses exhibit prolonged doubling times. Integrating phylogenetic eigenvectors into the ecological feature matrix, we employed a phylogenetic random forest model to identify key ecological features influencing defense presence and abundance. Further phylogenetic regressions unveil nuanced dependencies of bacterial defenses on specific environmental factors, challenging assumptions of a universal defense system distribution and underscoring reliance on subtle ecological cues. Notably, symbiotic and endosymbiotic bacteria often exhibit reduced defense systems and negative correlations between defense system abundance and the minimal doubling time. Conversely, free-living and motile bacteria show significant positive correlations between minimal doubling time and defense system abundance. Moreover, we highlight the pivotal role of ecological variables like habitat specificity and nutrient availability in shaping bacterial growth rates and defense mechanisms. Our findings underscore the complexity of microbial interactions and stress the need to consider ecological context in understanding defense strategies. We propose that trade-offs between growth and defense are ubiquitous due to sporadically inefficient optimization of limited resources, particularly in populations with small effective sizes, where both mechanisms may weaken concurrently due to genetic drift. This challenges traditional notions of trade-offs and underscores the impact of ecological context on microbial strategies.

## Introduction

Bacterial viruses, also known as phages or bacteriophages, represent the most abundant biological entities on Earth. They significantly outnumber bacterial cells across various samples and microbiomes, with ratios reaching up to ∼15-25 fold, as observed in freshwater plankton (1, 2). In environments like seawater, this constant predation results in a substantial daily death rate of bacterioplankton, estimated between 25-40% (3, 4). To combat phage attacks, bacteria have evolved an arsenal of defense tactics targeting every stage of the phage’s life cycle. These tactics include diverse mechanistic approaches such as the restriction-modification (RM) system, the Clustered Regularly Interspaced Short Palindromic Repeat and associated protein (CRISPR-Cas) system, and the Abortive Infection (Abi) system (5-7), collectively referred to as antiviral defense systems.

Despite the potential for horizontal gene transfer, bacteria typically only possess a subset of the available anti-phage mechanisms (8). For instance, the CRISPR-Cas systems, which provide adaptive immunity by targeting previously encountered pathogens, are found in approximately 40% of bacteria (9). Similarly, Abi systems are sporadically distributed across bacterial phylogeny (7), while RM systems are widespread but absent in about 4% of bacterial genomes (10). Additionally, the uneven distribution of antiviral defense systems extends to the archaeal lineage (9, 10), reflecting trade-offs between the costs of maintaining such systems and the benefits gained in phage resistance (8, 11-15).

In biological systems, traits often exhibit trade-offs due to constraints on resources such as energy, habitat, or time (16-18). For example, energy allocated for growth cannot simultaneously be used for defense or other energy-intensive processes, leading to necessary compromises among traits. While the concept of growth-defense trade-offs is well-established in plants and animals, its relevance in virus-infected prokaryotes is less recognized (11, 19).

Laboratory research on phage resistance has extensively explored trade-offs using model bacteria and phages such as *Escherichia coli* and *Pseudomonas aeruginosa*. Phage resistance evolution significantly impacts bacterial fitness components, including slowed cell division rates, reduced nutrient uptake, decreased virulence, and impaired cell motility (11, 20-29). Notably, some phage-resistant strains exhibit trade-ups, where selection improves multiple fitness components simultaneously, including antibiotic resistance (30, 31).

However, resistance to phages does not consistently correlate negatively with bacterial growth rates in natural environments. Studies examining marine *Synechococcus* strains and phage isolates, as well as bacteria and phages from horse chestnut leaves, have shown conflicting results regarding the growth-defense trade-off (26, 32). Recent research on marine bacterial communities has observed a negative correlation between growth rates and resistance to protists and viruses, supporting the growth-defense trade-off hypothesis (33).

To gain a comprehensive understanding of the trade-off between prokaryotic growth and defense, investigations encompassing diverse environments and species are essential. Recent research analyzing growth rates and adaptive immunity across bacterial domains revealed a significant positive correlation, indicating evolutionary trade-offs between growth and adaptive defense against virulent phages (15). Additionally, besides CRISPR-Cas systems, prokaryotes possess numerous other antiviral mechanisms, such as the RM and Abi systems, warranting further exploration (5, 6, 34, 35). Understanding the relationship between growth and antiviral defense across different environments and prokaryotic lineages holds promise for advancing our knowledge of evolution and informing the development of antimicrobial strategies.

To grasp the trade-off between prokaryotic growth and defense, it is crucial to take a comprehensive approach. Despite occasional discrepancies, our objective is to achieve a comprehensive understanding of whether phage-resistant lineages typically demonstrate slower growth rates compared to phage-sensitive ones across bacterial domains. These insights could significantly impact our understanding of evolutionary processes and contribute to the development of innovative antimicrobial strategies.

Recently, we investigated the relationship between growth rate and adaptive immunity in 4142 bacteria, utilizing predicted minimal doubling times retrieved from the EGGO database and CRISPR-Cas systems annotated with CRISPRCasFinder (15, 36, 37). Our findings unveiled a significant positive correlation, indicating evolutionary trade-offs between growth and adaptive defense against virulent phages within the bacterial domain.

In addition to CRISPR-Cas systems, prokaryotes have evolved various other antiviral mechanisms, including the RM system, the Abi system, and numerous others that warrant further exploration (5, 6, 34, 35). Notably, certain prokaryotes, such as the delta proteobacteria *Desulfonema limicola*, can harbor over 50 defense systems in their genomes (38). Recently, Tesson et al. (38) developed DefenseFinder, a novel antivirus-system detection program based on the HMM mapping algorithm, which enables the identification of known antiviral systems across 21,000 fully sequenced prokaryotic genomes. Leveraging this tool, we explored the relationship between growth and antiviral defense across bacterial domains, as well as within specific environments.

## Materials and Methods

The minimal doubling times of bacteria and archaea were extracted from the EGGO database (accessed on 2021-11-26), encompassing over 200,000 predictions of estimated growth rates (36). These values underwent a log transformation using the natural logarithm as the base. Due to its smaller sample size, we opted not to utilize the dataset of prokaryotic empirical minimal doubling times provided by Madin et al. (39) in this study. While empirical data typically offer higher accuracy compared to computationally generated predictions, predictive methodologies have their advantages (15). Empirical minimal doubling periods may offer superior advantages over computationally generated predictions due to the inherent limitations of predictive methodologies achieving 100% precision. However, the wide spectrum of genetic diversity observed among strains of individual bacterial species necessitates empirical data encompassing various factors, all procured from equivalent strains, a requirement that is not consistently met. Predicted phenotype values, including defense systems and growth rate, for a strain originate from the genome sequences of that very strain, potentially circumventing faults stemming from polymorphism within each species.

The antiviral defense systems of over 21,000 complete microbial genomes were sourced from Tesson et al. (38). Phylogenetic relationships among the analyzed prokaryotes were obtained from the Genome Taxonomy Database (GTDB) (40) (accessed 2022-04-08). GTDB offers superior taxonomic resolution and more precise classification compared to traditional phylogenetic trees by leveraging genomic data.

The intersecting samples among the three datasets mentioned above consisted of 3532 bacterial genomes and 237 archaeal genomes, some of which belonged to the same species. However, upon intersecting with subsequent datasets, only a minimal number of archaeal species were acquired, rendering most analyses impractical. Consequently, this paper focuses solely on the study of bacteria.

Ecological characteristics of prokaryotes were obtained from the ProTraits database (41), accessed on October 2, 2022, via “ProTraits_precisionScores.txt”. This comprehensive database integrates data from various repositories such as GOLD, NCBI, and KEGG, utilizing automated text mining and comparative genomic techniques to associate microorganisms with phenotypes. Our study focuses on ecological features potentially influencing defense system distribution, excluding physiological traits like glycerol and glutamate. Additionally, features with ambiguous biological significance, automated physiological features from machine learning, and those with missing *P* values exceeding 10% of the overall sample were discarded, retaining 125 prominent ecological features for analysis. Two columns represent each ecological feature in the database, providing precision scores for both the presence and absence of the feature, respectively. To facilitate analysis, these scores were converted into an overall maximal confidence score ranging from 0 to 1, following the methodology of Weissman et al. (42). Missing values in the ProTraits database were imputed using phylogenetic information (43). The PVRdecomp function of the R package PVR (Version 0.3) (44) was employed to extract eigenvectors from the phylogenetic distance matrix. Subsequently, the missForest function of the R package missForest (Version 1.5) (45) was utilized to estimate missing values.

Random forest, an integrated algorithm of decision trees, was utilized for regression tasks in this study. This method optimizes regression tasks, effectively handles high dimensionality, mitigates overfitting risks, accommodates both categorical and numerical variables, and consistently delivers reliable outcomes, making it superior to alternative techniques. The eigenvectors extracted from the phylogenetic distance matrix were incorporated into the ecological feature matrix to control phylogenetic relationships within the random forest analysis. The randomForest package (version 4.7-1.1) (46) was implemented in R (version 4.2.0).

Phylogenetic generalized least-squares (PGLS) analysis of correlations was conducted using the R package phylolm (version 2.6.2) (47). The Pagel’s λ model was employed due to its generally lower AIC values in our dataset. Differences in minimal doubling times between the two groups were compared using the phylANOVA function (48), while phylogenetic logistic regression analysis with dichotomous variables as dependent variables was performed using the phyloglm function (49). When multiple correlation analyses were performed on the same data, *P* values were corrected using the Benjamini & Hochberg method via the p.adjust function in R (Version 4.0.2).

## Results

### Phylogenetic Signals in Analyzed Features and Their Implications for Comparative Analysis

Due to shared ancestry among bacterial samples, feature values may exhibit non-independence, violating fundamental assumptions of conventional statistical methods. As such, using phylogenetic comparative methods becomes imperative to mitigate these effects (50, 51). We evaluated the phylogenetic signals of all analyzed features and observed significant signals in nearly all cases, with minimal doubling time demonstrating the strongest signal (Supplementary Table S1). Consequently, we will employ phylogenetic comparative methods whenever applicable.

A recent study by Chen et al. (52) highlighted the potential for conflicting outcomes in PGLS analysis when interchanging dependent and independent variables. While PGLS cannot establish causality but rather identifies correlations, Chen et al. utilized simulated data and a gold standard to establish criteria for selecting the dependent variable. Their findings suggested that choosing the feature with a stronger phylogenetic signal as the dependent variable led to more accurate correlation detection than randomly selected variables. In line with this approach, we adopted the feature exhibiting a stronger phylogenetic signal as the dependent variable in all PGLS analyses conducted in this study.

### Ecological Factors Influencing the Presence of Antiviral Defense Systems in Bacteria

Our analysis stems from a comprehensive program capable of detecting all known antiviral systems in prokaryotic genomes (38). However, among the 3532 bacterial genomes (Supplementary Table S2) examined, 7% (243) were found to lack any defense systems. Phylogenetic ANOVA analysis unveiled that these bacteria exhibited significantly longer minimal doubling times compared to those encoding one or more defense systems (Mean ± SD: 5.6 ± 3.1 h *vs*. 3.2 ± 4.4 h. *P* = 0.0011). Surprisingly, bacteria devoid of defense systems did not demonstrate faster proliferation rates, contradicting the anticipated trade-off between growth and defense. It is plausible that these bacteria inhabit specific ecological niches where neither defense mechanisms nor rapid growth are necessary.

To delve into the factors influencing the presence or absence of bacterial defense systems, we leveraged ecological data from the ProTraits database. Since the database primarily comprises bacterial species rather than individual strains or genomes, we aggregated some of the 3532 genomes into species. To capture species-level characteristics, we computed median values of defense system abundance and minimal doubling time for genomes within the same species, identifying 1044 distinct bacterial species (Supplementary Table S3).

Utilizing a random forest model, we ranked the importance of 125 ecological features. Remarkably, factors such as “known habitats = insect endosymbiont,” “known habitats = endosymbiont,” “biotic relationship = symbiont,” and “host = insects general” emerged as top influencers, underscoring the significance of symbiotic and endosymbiotic relationships (Fig. 1).

**Figure 1.**
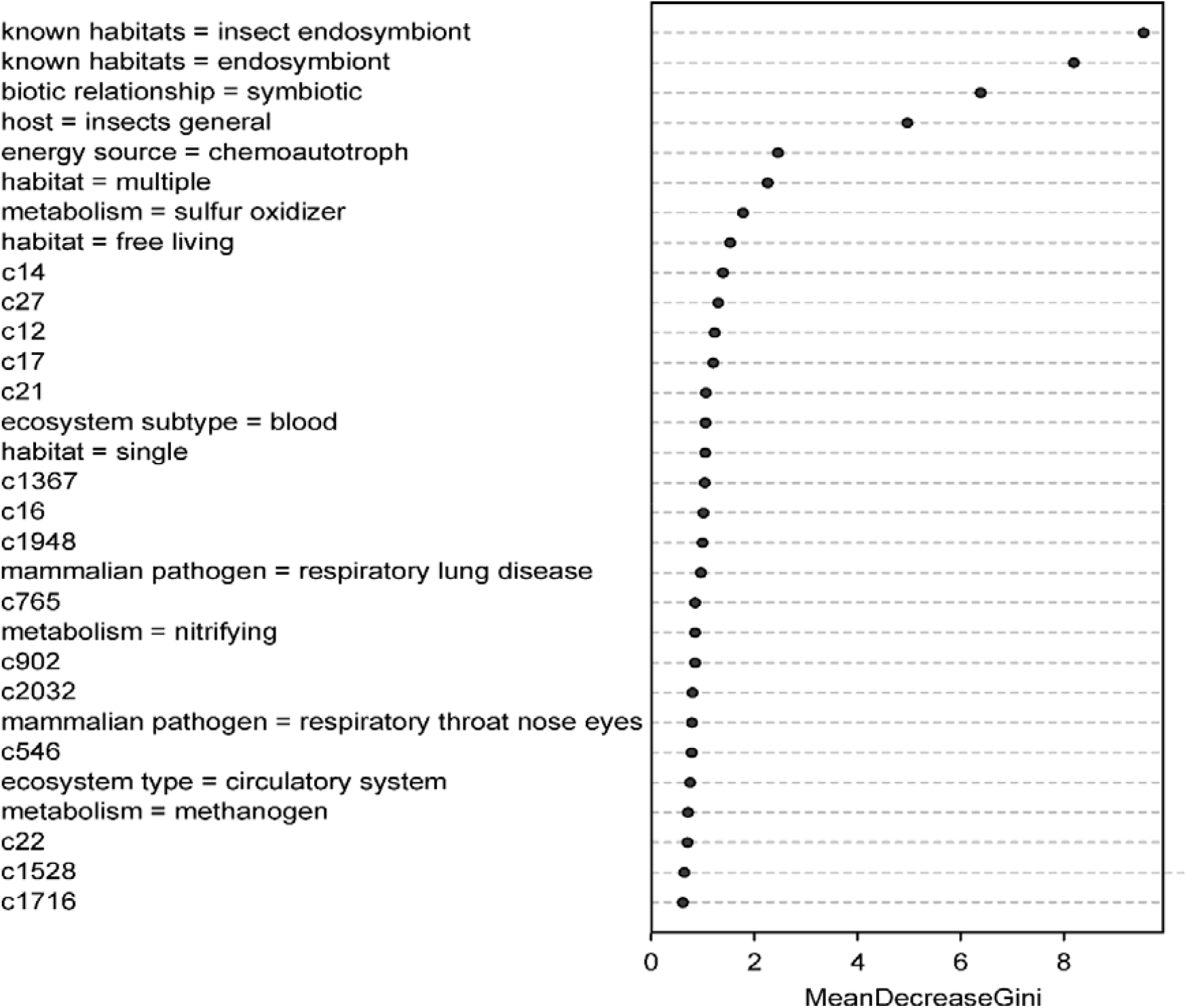
Importance ranking of ecological features for defense system presence/absence in bacteria. The vertical axis represents the names of ecological features, with higher ranks indicating a greater impact on the presence or absence of the defense system. MeanDecreaseGini, displayed on the horizontal axis, quantifies feature importance, with higher values indicating greater significance. The figure displays the top thirty features in the model. Features starting with the letter “c” represent eigenvectors representing phylogenetic relationships.

Given that the random forest model cannot discern the directional effect of features on the presence or absence of antiviral defense systems, we conducted phylogenetic logistic regression (Table 1). Results indicated a negative correlation between antiviral defense presence and “known habitats = insect endosymbiont,” “biotic relationship = symbiont,” and “host = insects general” (*P* < 10^−8^ for all), while “known habitats = endosymbiont” showed no significant correlation (*P* = 0.452).

**Table 1.** Phylogenetic logistic regression of defense system presence/absence on bacterial habitats.

| Dependent variable ~ independent variable | Slope | *P* | *P*<sub>adjusted</sub> |
| --- | --- | --- | --- |
| Presence/absence of defense ~ Known habitats = insect endosymbiont | -16.108 | $5.6 \times 10^{-16}$ | $2.3 \times 10^{-15}$ |
| Presence/absence of defense ~ Known habitats = endosymbiont | 0.215 | 0.543 | 0.543 |
| Presence/absence of defense ~ Biotic relationship = symbiotic | -7.387 | $6.6 \times 10^{-10}$ | $1.3 \times 10^{-9}$ |
| Presence/absence of defense ~ Host = insects general | -8.558 | $3.4 \times 10^{-9}$ | $4.5 \times 10^{-9}$ |

The default parameter logistic MPLE was used in the phylogenetic logistic regression of the relationships. In the regression analysis, the presence of defense systems was assigned as 1 and the absence was assigned as 0. The habitats are measured by the overall maximal confidence score described in Materials and Methods section. The ratio of samples with defense systems to those without related systems was 965 vs 79. The *P*<sub>adjusted</sub> value represents the corrected *P*-value computed using the Benjamini & Hochberg method.

### Ecological Determinants of Defense System Abundance: Insights from Symbiosis and Environmental Dynamics

The preceding study explored factors influencing the presence or absence of defense systems. Here, we delve deeper into the variables shaping defense system abundance. Bacteria lacking defense mechanisms will be recorded as having zero systems, ensuring inclusivity in our analysis.

Among 125 ecological features, “known habitats = insect endosymbiont,” “known habitats = endosymbiont,” and “biotic relationship = symbiont” emerged as top influencers, underscoring the importance of symbiotic associations (Fig. 2). Furthermore, symbiotic relationships with hosts were supported by additional features within the top rankings, including “ecosystem type = circulatory system,” “ecosystem subtype = blood,” “habitat = host associated,” “host = insects general,” “ecosystem category = mammals,” “known habitats = host,” and “ecosystem = host associated.” If symbiotic bacteria tend to have fewer defense systems for certain reasons, free-living bacteria should exhibit a comparatively higher abundance of defense systems. Indeed, “habitat = free living” ranks within the top 19.

**Figure 2.**
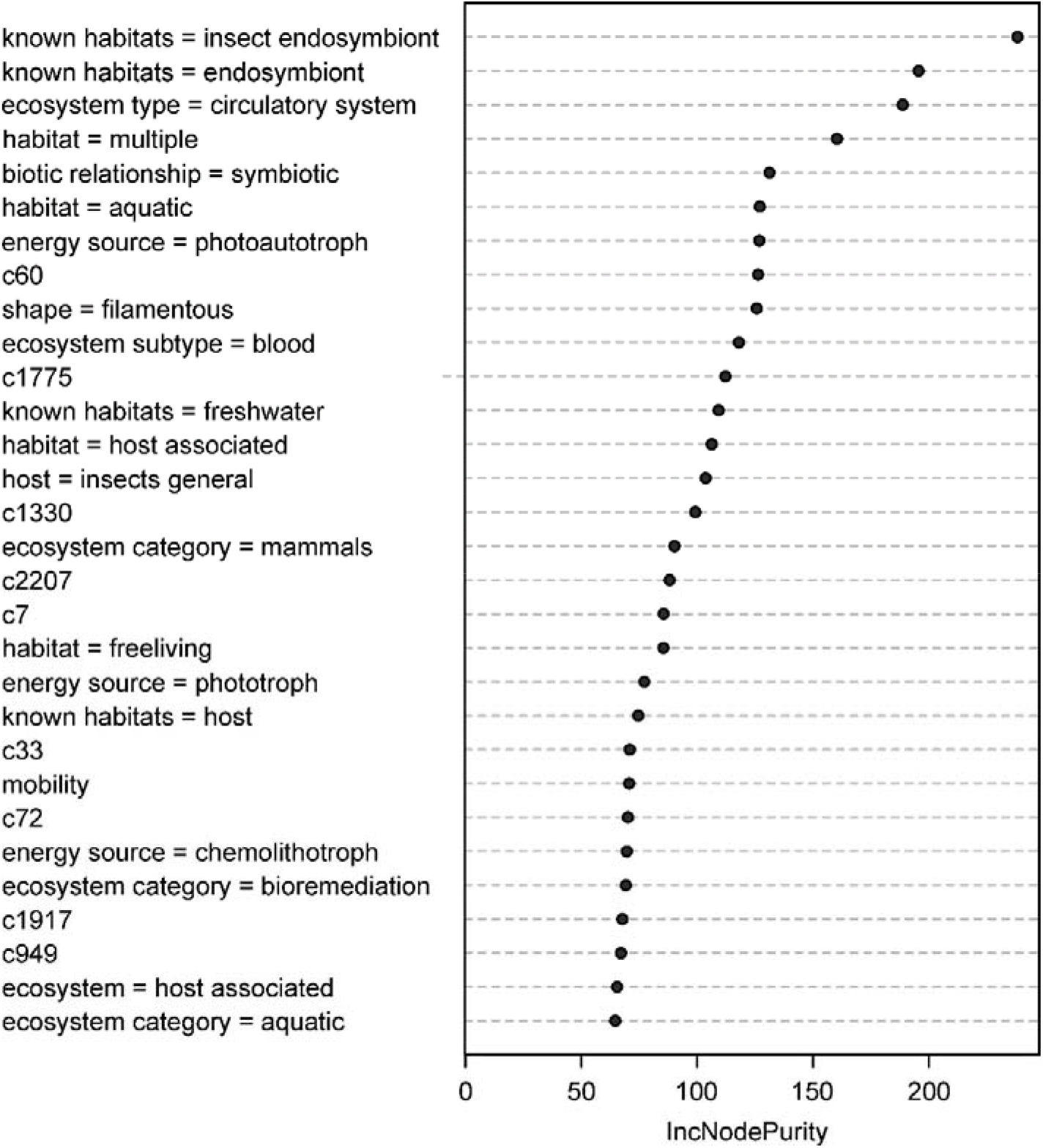
Importance ranking of ecological features for defense system abundance in bacteria. The vertical axis represents the names of ecological features, with higher ranks indicating a greater impact on the abundance of the defense system. IncNodePurity, displayed on the horizontal axis, quantifies feature importance, with higher values indicating greater significance. The figure displays the top thirty features in the model. Features starting with the letter “c” represent eigenvectors representing phylogenetic relationships.

Additionally, aquatic environments and energy sources emerged as key factors. “Habitat = aquatic” and “energy source = photoautotroph” ranked prominently, indicating their influence on defense system abundance (Fig. 2). Additionally, polyhabitat, filamentous shape, mobility, and bioremediation also appear to influence defense system abundance significantly.

Further PGLS analysis revealed negative correlations between symbiotic characteristics and defense system abundance, while free-living bacteria exhibited a positive association (Table 2). While some correlations may not achieve statistical significance, all other ecological characteristics positively correlate with the abundance of defense systems (Table 2). Notably, bacteria inhabiting multiple habitats, exhibiting motility, or being involved in environmental bioremediation tend to possess a higher abundance of defense systems. Given their exposure to complex or fluctuating environments, these bacteria are more susceptible to encountering invasive elements such as phages. Another intriguing finding is that filamentous bacteria harbor significantly more defense systems, likely due to their filamentous morphology providing increased sites for phage attachment.

**Table 2.** PGLS regressions of ecological features on the defense system abundance in bacteria.

|  | Slope | *P* | *P*<sub>adjusted</sub> |
| --- | --- | --- | --- |
| Known habitats = insect endosymbiont | -0.001 | 0.001 | 0.004 |
| Known habitats = endosymbiont | -0.001 | $4.4 \times 10^{-4}$ | 0.002 |
| Biotic relationship = symbiotic | -0.002 | 0.022 | 0.031 |
| Ecosystem type = circulatory system | -0.001 | 0.019 | 0.029 |
| Ecosystem subtype = blood | -0.001 | 0.068 | 0.084 |
| Habitat = host associated | -0.002 | 0.002 | 0.005 |
| Host = insects general | -0.002 | 0.002 | 0.005 |
| Ecosystem category = mammals | -0.002 | 0.015 | 0.024 |
| Known habitats = host | -0.002 | 0.008 | 0.017 |
| Ecosystem = host associated | -0.002 | 0.011 | 0.019 |
| Habitat = free living | 0.002 | 0.007 | 0.016 |
| Habitat = aquatic | 0.001 | 0.054 | 0.071 |
| Known habitats = freshwater | 0.001 | 0.190 | 0.210 |
| Ecosystem category = aquatic | 0.001 | 0.284 | 0.284 |
| Energy source = photoautotroph | 0.001 | 0.251 | 0.264 |
| Energy source = phototroph | 0.003 | $8.5 \times 10^{-6}$ | $9.0 \times 10^{-5}$ |
| Energy source = chemolithotroph | 0.001 | 0.105 | 0.123 |
| Habitat = multiple | 0.002 | 0.001 | 0.004 |
| Shape = filamentous | 0.002 | $3.8 \times 10^{-6}$ | $8.0 \times 10^{-5}$ |
| Mobility | 0.002 | 0.010 | 0.019 |
| Ecosystem category = bioremediation | 0.002 | $3.9 \times 10^{-5}$ | $2.7 \times 10^{-4}$ |

### Ecological Factors Impacting Bacterial Growth Rates: Insights from Endosymbiosis and Habitat Dynamics

We investigate whether ecological factors influencing defense system abundance affect bacterial growth rates. Using PGLS regression, we correlated minimal doubling time with overall maximal confidence scores for significant ecological features.

Endosymbiosis and symbiosis with insects differ in their effects on bacterial growth rates compared to symbiosis with mammals (Table 3). “Known habitats = insect endosymbiont,” “Known habitats = endosymbiont,” and “Host = insects general” exhibit significantly positive correlations with minimal doubling time. Conversely, features related to symbiosis with mammals show negative correlations, albeit not statistically significant. Ambiguous features like “Biotic relationship = symbiotic” show a significant positive correlation, while others like “Ecosystem = host associated,” “Known habitats = host,” and “Habitat = host associated” exhibit negative correlations.

**Table 3.**
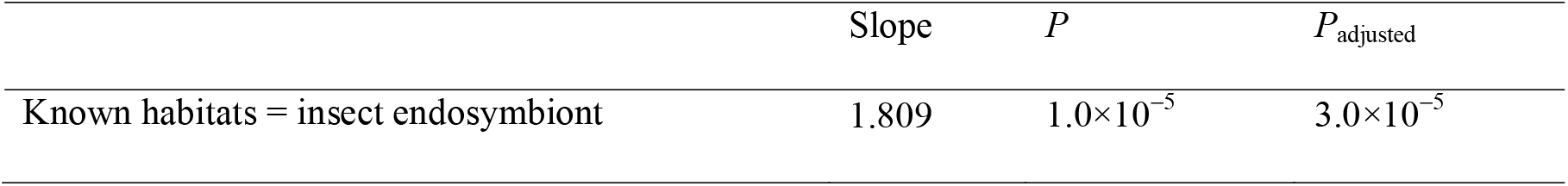
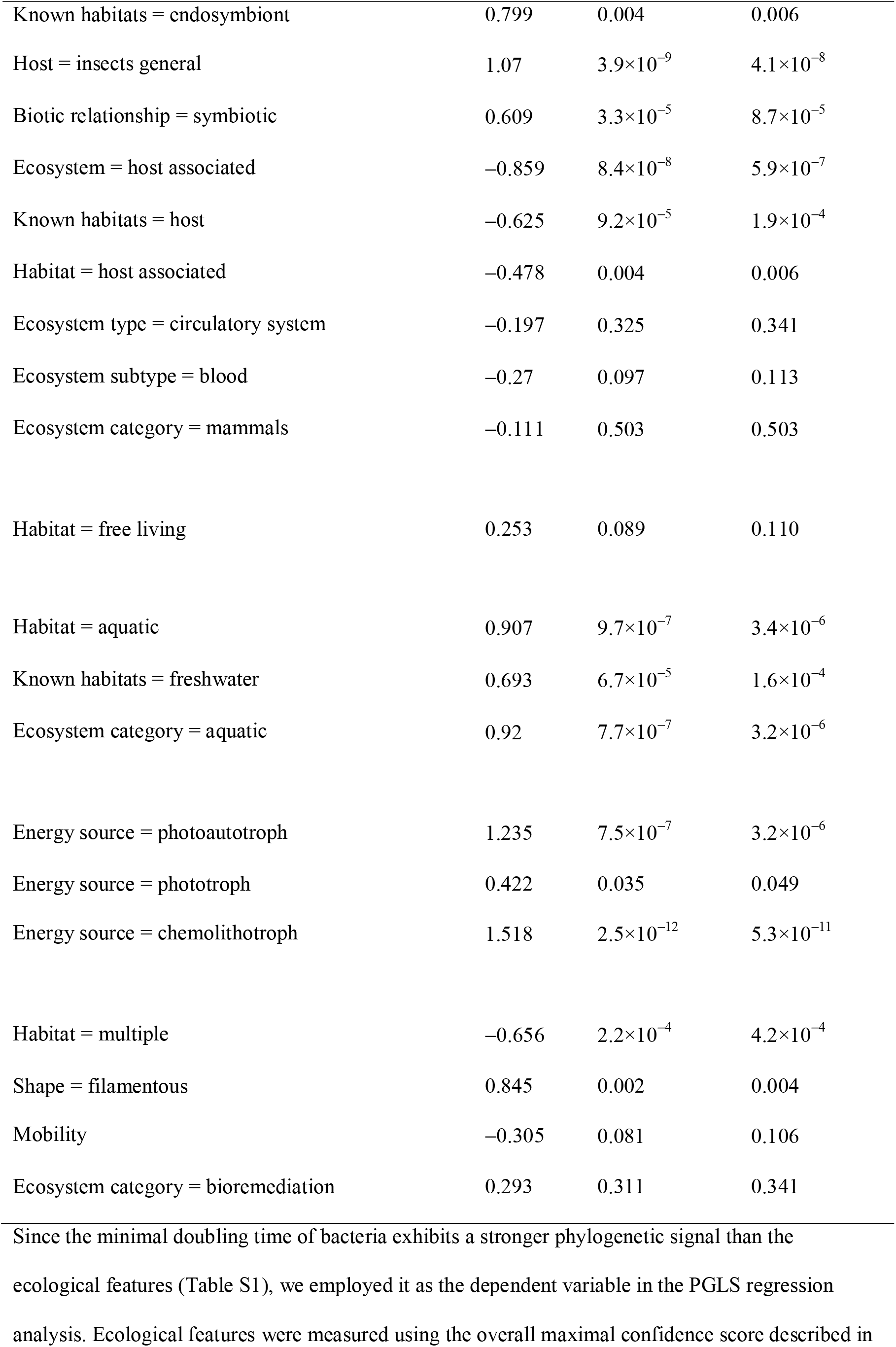
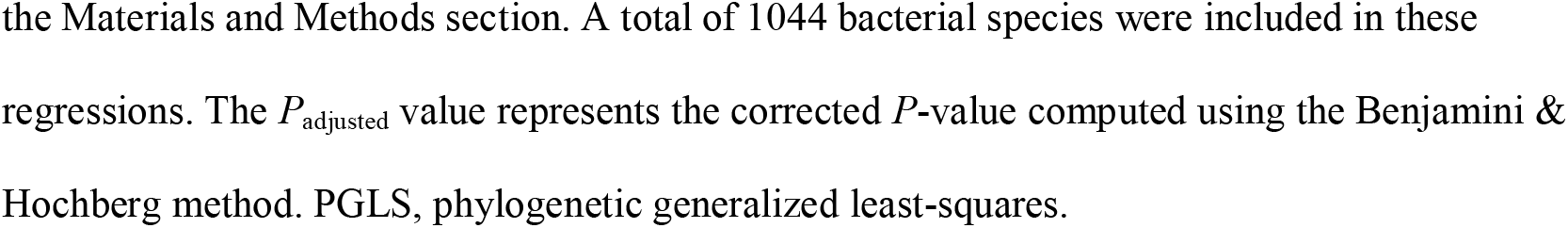
PGLS regressions of bacterial minimal doubling time on ecological features.

| | Slope | $P$ | $P_{\text{adjusted}}$ |
| --- | --- | --- | --- |
| Known habitats = insect endosymbiont | 1.809 | $1.0 \times 10^{-5}$ | $3.0 \times 10^{-5}$ |
| Known habitats = endosymbiont | 0.799 | 0.004 | 0.006 |
| Host = insects general | 1.07 | $3.9 \times 10^{-9}$ | $4.1 \times 10^{-8}$ |
| Biotic relationship = symbiotic | 0.609 | $3.3 \times 10^{-5}$ | $8.7 \times 10^{-5}$ |
| Ecosystem = host associated | -0.859 | $8.4 \times 10^{-8}$ | $5.9 \times 10^{-7}$ |
| Known habitats = host | -0.625 | $9.2 \times 10^{-5}$ | $1.9 \times 10^{-4}$ |
| Habitat = host associated | -0.478 | 0.004 | 0.006 |
| Ecosystem type = circulatory system | -0.197 | 0.325 | 0.341 |
| Ecosystem subtype = blood | -0.27 | 0.097 | 0.113 |
| Ecosystem category = mammals | -0.111 | 0.503 | 0.503 |
| Habitat = free living | 0.253 | 0.089 | 0.110 |
| Habitat = aquatic | 0.907 | $9.7 \times 10^{-7}$ | $3.4 \times 10^{-6}$ |
| Known habitats = freshwater | 0.693 | $6.7 \times 10^{-5}$ | $1.6 \times 10^{-4}$ |
| Ecosystem category = aquatic | 0.92 | $7.7 \times 10^{-7}$ | $3.2 \times 10^{-6}$ |
| Energy source = photoautotroph | 1.235 | $7.5 \times 10^{-7}$ | $3.2 \times 10^{-6}$ |
| Energy source = phototroph | 0.422 | 0.035 | 0.049 |
| Energy source = chemolithotroph | 1.518 | $2.5 \times 10^{-12}$ | $5.3 \times 10^{-11}$ |
| Habitat = multiple | -0.656 | $2.2 \times 10^{-4}$ | $4.2 \times 10^{-4}$ |
| Shape = filamentous | 0.845 | 0.002 | 0.004 |
| Mobility | -0.305 | 0.081 | 0.106 |
| Ecosystem category = bioremediation | 0.293 | 0.311 | 0.341 |

the Materials and Methods section. A total of 1044 bacterial species were included in these regressions. The $P_{\text{adjusted}}$ value represents the corrected $P$ -value computed using the Benjamini & Hochberg method. PGLS, phylogenetic generalized least-squares.

Aquatic habitat and energy source features positively correlate with minimal doubling time (Table 3), suggesting slower growth rates in bacteria residing in aquatic environments and utilizing natural energy sources. Additionally, filamentous morphology correlates with slower growth rates, while bacteria inhabiting multiple habitats tend to grow faster (Table 3).

### Exploring Growth-Defense Relationships Across Ecological Niches

In our analysis utilizing the ProTraits database, each species received an overall maximal confidence score indicating the likelihood of possessing specific ecological features. We categorized species into presence or absence groups based on a threshold of 0.5, compiling species counts for the top 30 influential ecological features affecting defense system abundance. Features with fewer than 20 species were filtered out.

To investigate potential growth-defense trade-offs within these ecological niches, we conducted PGLS regression analysis, correlating bacterial minimal doubling time with defense system abundance (Table 4). Highly significant correlations (*P* < 0.01) were primarily observed in two niches: “Habitat = free living” and “Mobility,” with extensive sample sizes comprising hundreds of bacterial species.

**Table 4.** PGLS regressions of bacterial minimal doubling time on the abundance of defense systems.

|  | *n* | Slope | *P* |
| --- | --- | --- | --- |
| Host = insects general | 30 | 0.092 | 0.478 |
| Biotic relationship = symbiotic | 98 | −0.009 | 0.210 |
| Ecosystem = host associated | 499 | −0.015 | 0.028 |
| Known habitats = host | 518 | −0.016 | 0.021 |
| Habitat = host associated | 434 | −0.017 | 0.020 |
| Ecosystem type = circulatory system | 52 | −0.012 | 0.194 |
| Ecosystem subtype = blood | 92 | −0.018 | 0.104 |
| Ecosystem category = mammals | 81 | −0.019 | 0.129 |
| Habitat = free living | 605 | 0.013 | 0.008 |
| Habitat = aquatic | 194 | 0.007 | 0.265 |
| Known habitats = freshwater | 147 | 0.001 | 0.901 |
| Ecosystem category = aquatic | 302 | 0.007 | 0.243 |
| Energy source = photoautotroph | 20 | 0.028 | 0.023 |
| Energy source = chemolithotroph | 22 | 0.01 | 0.524 |
| Mobility | 410 | 0.02 | 0.002 |
| Habitat = multiple | 206 | 0.01 | 0.264 |
| Shape = filamentous | 22 | −0.001 | 0.837 |

Among the features related to symbiosis, significant correlations between growth and defense were identified in three features, each representing samples of over 400 species: “Ecosystem = host associated,” “Known habitats = host,” and “Habitat = host associated.” Accounting for phylogenetic data, closely related lineages with similar characteristics were treated as nearly one effective sample. Thus, the effective sample size in phylogenetic comparative analysis was substantially lower than the total number of analyzed lineages. Consequently, the conventional statistical rule of thumb regarding small and large samples (*n* = 30) should not be applied in phylogenetic comparative studies (53).

Nevertheless, the findings presented in Table 4 suggest the presence of not only a trade-off but also a trade-up between bacterial growth and defense.

### Contextual Dependency in Growth-Defense Correlations Across Bacterial Environments

In Table 4, diverse correlations emerge between bacterial growth and defense across different environments. Positive correlations are observed in some bacteria within specific habitats, while negative correlations are apparent in others; meanwhile, no correlation is discernible in some instances. Consequently, within a large sample, the prevalence of each bacterial type dictates the observed correlation pattern: a predominance of bacteria exhibiting positive correlations yields a positive overall correlation, a balance of the three types results in no discernible correlation, and a prevalence of bacteria with negative correlations leads to an overall negative correlation. Thus, the significance of analyzing growth-defense correlations on a global scale diminishes.

Nevertheless, we conducted PGLS regression analysis to explore the growth-defense correlation in our dataset of 3532 samples, yielding statistically insignificant results (*P* = 0.398). Despite the lack of statistical significance, our analysis underscores the complex and context-dependent nature of the relationship between bacterial growth rates and defense mechanisms across diverse ecological contexts.

## Discussion

The analysis of bacterial defense mechanisms, whether assessed by the presence or absence of defense systems, or their abundance, yielded notable findings: ecological factors associated with symbiosis and endosymbiosis emerged as the primary influencers of bacterial defenses (Figs. 1, 2). Furthermore, phylogenetic regression analyses demonstrated a trend wherein symbiotic or endosymbiotic bacteria tend to exhibit fewer defense systems or lack them altogether (Table 1, 2). This observation aligns with prior findings noting the scarcity of defense systems among insect endosymbionts (38, 54-60). Our study unveils statistically significant associations between symbiosis/endosymbiosis and defense mechanisms, suggesting that such bacteria have reduced the necessity for robust defensive strategies. Endosymbiotic bacteria, residing within host cells, enjoy physical insulation from aggressive phages. Similarly, evidence suggests that many symbiotic bacteria are spatially separated from such phages. Notably, the gut microbiome represents a well-studied example of symbiotic bacteria, comprising the majority of human symbionts (61, 62). Within the mammalian small intestine and colon, specialized epithelial cells secrete a mucosal layer (63), which serves as a formidable barrier against the spread and access of phage populations, providing a refuge for bacteria (64-66). Consequently, the phage-bacterial ratios in the human gut are significantly lower compared to other ecosystems, often ranging from 1:1 to 0.1:1 as opposed to 10:1(67). In addition, the low pH in the stomach, digestive enzymes (mainly proteases), and bile salts make the gastrointestinal tract hostile to phages (68-70), contributing to a lower virus-to-bacteria ratio in the gut compared to other environments.

However, the impact of endosymbiosis on bacterial growth rates in insects differs significantly from symbiosis in mammals (Table 3). The proliferation of endosymbionts is not solely determined by their inherent ability to replicate but is also influenced by regulatory constraints imposed by their hosts (71, 72). In contrast, the gut presents a highly complex environment for bacteria. Nutrient levels fluctuate considerably on a daily basis, and the presence of antimicrobial molecules and oxygen secreted by the epithelium strongly restricts potential microbial inhabitants (63), creating a gradient of suitability for bacterial growth from the lumen to the mucosa. Bacteria residing in isolated microhabitats such as crypts may be tightly controlled by the host epithelium, while those dispersed into the gut lumen could proliferate in response to nutrient availability. We hypothesize that the overall impact of gut symbiosis on bacterial growth rate depends on the distribution of bacteria across different microhabitats.

Due to insufficient sample size, no statistical results have been yielded regarding the relationship between growth and defense in endosymbiotic bacteria. However, a weak negative correlation between minimal doubling time and the abundance of defense systems was observed in symbiotic bacteria (Table 4). This observation suggests that symbiotic bacteria with robust defense mechanisms tend to exhibit rapid growth. We speculate that, despite potential fluctuations in nutrient levels, hosts typically provide a eutrophic environment for symbiotic bacteria, where energy and resources are not limiting factors, thus rendering the trade-off between growth and defense unnecessary (18). The gut lumen may favor *r*-selected bacteria, which prioritize rapid growth and reproduction in unstable environments.

Several ecological features, such as free-living habitat, phototrophic lifestyle, occupancy of multiple habitats, motility, filamentous morphology, and participation in bioremediation, have been positively linked to the abundance of defense systems (Table 2). These findings expand upon previous research on the relationship between defense system abundance and lifestyle, habitat, and geographic background (38, 54-60). The trade-off between growth and defense has been statistically demonstrated in free-living and motile bacteria. These findings suggest that certain ecological niches may favor the allocation of resources towards defense mechanisms, potentially at the expense of growth rates, in order to cope with environmental challenges such as predation by bacteriophages. The positive association between defense system abundance and ecological features like free-living habitat and motility implies that bacteria thriving in diverse and dynamic environments may require heightened defense strategies to survive and proliferate.

Previous studies have shown a strong association between the abundance of defense systems and genome size, particularly noting the absence of defense genes in the small genomes of endosymbionts (38, 56, 60, 73-75). In fact, one study suggests that the number of defense systems remains largely unaffected by the lifestyle of microbes after controlling for genome size, attributing the scarcity of defense genes in endosymbionts to their small genomes rather than their lifestyle (60). In our view, while there is a significant correlation between genome size and the abundance of defense systems, it does not necessarily mean that studying the quantity of defense systems makes sense only after controlling for genome size. If, for some reason, the evolutionary driving force behind an increase in genome size leads to an increase in defense systems, or conversely, if the evolutionary driving force behind a decrease in genome size leads to a reduction in defense system genes, then conducting partial correlation analysis controlling for genome size would be necessary. However, the reduction in genome size in endosymbionts is not a result of natural selection but rather a consequence of stochastic drift; the genome size decreases because genes are lost. The small genomes of endosymbionts are the result of many gene losses, including defense systems, rather than a parallel phenomenon to gene loss. Therefore, controlling for genome size, which effectively eliminates the gene loss itself, is not only meaningless but also misleading when studying the evolution of the quantity of defense system genes (both gained and lost).

The concept of trade-offs in evolutionary biology underscores how organisms adapt their life strategies in response to limited resources (17, 18). While certain environments clearly necessitate a trade-off between growth and defense strategies, in others, this distinction may become less evident. Firstly, the resource may occasionally unlimited, the requirement of both growth and defense could be simultaneously met. Secondly, and more commonly, this ambiguity may arise due to weak selection efficiency or the prevalence of stochastic drift, especially noticeable in populations with small effective sizes, which can hinder optimization. In this case, resources saved in one aspect may not be efficiently reallocated to another. Given this perspective, some endosymbiotic and symbiotic bacteria might grow slowly despite lacking robust defense systems, challenge the traditional notion of a positive correlation between growth and defense. Labeling this phenomenon as a trade-up, as previously done to describe inconsistencies with conventional trade-offs (23, 76-78), may not suffice. Instead, with smaller effective population size and less efficient natural selection (79), both growth and defense mechanisms may weaken simultaneously, suggesting that it might be more accurately termed a trade-down. Expanding on this line of thought, we find an explanation for the dependence of growth and defense on ecological contexts. In various environments, the majority of bacterial species may exhibit differing effective population sizes. In environments where most bacterial species boast large effective population sizes, indicating high natural selection efficiency, a trade-off between growth and defense, resulting in a negative correlation, is likely to occur. However, in habitats dominated by species with small populations, where genetic drift prevails, life strategies cannot be optimized, thereby nullifying the trade-off between growth and defense.

## Supplementary Information

Supplementary file 1 in docx format: Table S1. Phylogenetic signals of the bacterial features analyzed in this study.

Supplementary file 2 in xlsx format: Table S2. A comprehensive dataset of the minimal doubling time and antiviral defense system features for 3532 bacterial genomes. Table S3. A comprehensive dataset of ecological characteristics, phylogenetic feature vectors, antiviral defense systems, and other features for 1044 bacterial species.

## Acknowledgments

We are grateful to Tian-Hao Zhao for his help in machine learning techniques and Quan-Guo Zhang for hekpful comments. This article has undergone refinement by ChatGPT 3.5 to enhance language and readability (80). Subsequently, the authors thoroughly reviewed the manuscript and made necessary edits, taking full responsibility for the content.

## Author contributions

Deng-Ke Niu conceived the research hypothesis; Zhi-Ling Liu and Jia Liu analyzed the data. Zhi-Ling Liu and Deng-Ke Niu drafted the manuscript; all authors contributed to the final version and reviewed the manuscript.

## Availability of data and materials

The datasets supporting the conclusions of this article are included within the article and its

Supplementary files.

## Declarations

### Ethics approval and consent to participate

All authors consent to participate in this study.

### Consent for publication

All authors give their consent for publication.

