## Supplemental Table 1 for "Bacterial defenses and their trade-off with growth are not ubiquitous but depend on ecological contexts"

Table S1. Phylogenetic signals of the bacterial features analyzed in this study

| Traits | *n* | Pagel’s λ | *P* |
| --- | --- | --- | --- |
| The abundance of defense systems | 1044 | 0.700 | 7.31×10^−62^ |
| Minimal doubling time | 1044 | 0.977 | 5.69×10^−259^ |
| Known habitats = insect endosymbiont | 1044 | 0.993 | 1.86×10^−274^ |
| Known habitats = endosymbiont | 1044 | 0.812 | 1.44×10^−122^ |
| Ecosystem type = circulatory system | 1044 | 0.872 | 3.19×10^−237^ |
| Habitat = multiple | 1044 | 0.906 | 6.66×10^−263^ |
| Biotic relationship = symbiotic | 1044 | 0.832 | 1.19×10^−180^ |
| Habitat = aquatic | 1044 | 0.925 | 2.10×10^−312^ |
| Energy source = photoautotroph | 1044 | 0.862 | 5.87×10^−290^ |
| Shape = filamentous | 1044 | 0.933 | 5.93×10^−323^ |
| Ecosystem subtype = blood | 1044 | 0.877 | 6.75×10^−257^ |
| Known habitats = freshwater | 1044 | 0.900 | 5.40×10^−270^ |
| Habitat = host associated | 1044 | 0.889 | 1.38×10^−270^ |
| Host = insects general | 1044 | 0.849 | 2.43×10^−111^ |
| Ecosystem category = mammals | 1044 | 0.841 | 2.76×10^−181^ |
| Habitat = freeliving | 1044 | 0.852 | 2.649×10^−232^ |
| Energy source = phototroph | 1044 | 0.820 | 1.457×10^−162^ |
| Known habitats = host | 1044 | 0.890 | 1.886×10^−260^ |
| Mobility | 1044 | 0.883 | 1.01×10^−287^ |
| Energy source = chemolithotroph | 1044 | 0.919 | 1.29×10^−261^ |
| Ecosystem category = bioremediation | 1044 | 0.826 | 2.35×10^−130^ |
| Ecosystem = host associated | 1044 | 0.910 | 2.85×10^−262^ |
| Ecosystem category = aquatic | 1044 | 0.922 | 1.26×10^−299^ |
